# Biomimetic Cues Enable Predictive Mechanisms in Simulated and Physical Robot-Human Object Handovers

**DOI:** 10.1101/2025.10.14.682495

**Authors:** Clara Günter, Yuhe Gong, Riddhiman Laha, Simon Appoltshauser, Luis Figueredo, Joachim Hermsdörfer, David W. Franklin

## Abstract

Object handovers – while representing one of the simplest forms of physical interaction between two agents – involve a complex interplay of predictive and reactive control mechanisms in both agents. As human-human pairs have unrivaled skills in physical collaboration tasks, we take the approach of understanding and applying biomimetic concepts to human-robot interaction. Here, we apply the concept of passer movement cues, that is, slower movement for heavy objects and faster movements for lighter objects, to robot-human handovers. We first show that when a simulated passing agent’s movement is scaled with object mass, participants as receivers adapt their anticipatory grip forces according to mass in a virtual environment. We then apply the same concept to a physical robot-human handover and show that our approach generalizes to the real-world. The predictive scaling of grip forces is learned iteratively upon repeated presentations of trajectory-mass pairings, whether the masses are presented in a random or blocked order. Overall we demonstrate that the presentation of robotic kinematic cues can provide intuitive and naturalistic human predictive control in object handover. This extends the use of non-verbal cues in robot-human handover tasks and facilitates more legible and efficient physical robot-human interactions.

## 1 Introduction

As robots become more immersed in humans’ everyday life, legibility, interpretability, and mutual understanding become paramount for achieving seamless interaction. This is particularly critical for foundational physical skills, which are expected to occur frequently and in diverse contexts. Among such skills, object handover stands out as both one of the most ubiquitous activities and deceptively complex [52].

The handover action itself can be divided into three main phases [14, 45]: (i) the passer transport phase, where the passing agent lifts and transports the object to the handover location, (ii) the physical handover, where both agents are in physical contact with the object, and (iii) the receiver transport phase, where the receiver transports the object and typically completes a secondary task.

The main phase used to measure efficiency and smoothness of the handover action is the physical handover (ii), where — in the ideal case — the passer releases the object in synchrony or minimal delay to the receiver grasping it. Remarkably, in human-human handovers, this delay has been observed to be faster than neural delays in humans [24], indicating the importance of predictive mechanisms. That is, human receivers must use such predictions to estimate e.g. object properties, before receiving physical feedback about them.

These anticipatory actions rely on predictive mechanisms informed by visual and contextual cues. Humans can learn to associate a wide variety of cues with object properties and scale interaction forces accordingly. Previous work has demonstrated the use of size [29], color [15], material [4], position [60] and even general familiarity [35], to predictively scale grip forces to object properties. While arbitrary (e.g. color) and contradictory (large object-light weight) cue-property pairs can be learned through experience [18], the adaptation process is extended and the strength of association reduced as compared to more familiar pairings [35].

In an object handover, beyond memorizing cues directly related to the object, observing the passing agent interact with the object may also inform the receiver about its properties. For example, when observing humans manipulating a novel object, unique movement characteristics are evident [16, 41]. Specifically, the delay between contact and liftoff for heavy objects tends to be longer [16, 41] and the transport peak velocity lower [41]. Consequently, such movement kinematics of the passer may be used as a cue by the receiver to predict object weight in human-human handovers [42]. However, existing studies lack the isolation of movement kinematics.Translating this predictive capability to robots requires understanding how humans interpret these signals, particularly when stemming from a human-robot interaction (HRI), and whether robots can embed them intentionally into their motion, particularly in close-proximity interactions where physical contact is imminent. In this context, while previous research has established a relationship between passer movement and receiver prediction in human–human handovers, several open challenges still remain. First, to the best of the authors’ knowledge, there is no study to evaluate if such effects transfer to robot–human interactions, as well as how the kinematic cue perception can be influenced by the robot motion. Furthermore, existing human-human studies cannot isolate the desired control variables from the passer’s own expectations, which, in turn, affect their behaviour. That is, studies have not fully disentangled the effect of prior expectations on movement kinematics, leaving a gap in biomechanical analysis, and in robotics, a gap in experimentation, transfer, evaluation, and method development. Addressing these questions is critical for advancing predictive, fluent, and safe physical collaboration between humans and robots.

## 2 Related Work

Object handover is a core skill in both human–human and human–robot collaboration, requiring precise coordination between two control systems on the same object in real time. Translating this fluid coordination to human–robot teams remains challenging [52]. Differences in motion style, predictability, and interaction dynamics can alter how cues are perceived, and prior work has highlighted that humans may behave differently when collaborating with robots compared to humans [6, 40, 53]. Understanding and evaluating such mechanisms and their transferability to robotic systems is therefore essential for improving robot handover fluency. In this context, this work focuses on two main areas, which have often been studied separately in the past, predictability during human-human handover and human-robot handover.

### 2.1 Predictability during Human-Human Handover

While predictability of object properties is important, general predictability of the partner’s actions in a joint task is essential for smooth collaboration [3, 11, 14, 38, 42]. Previous work demonstrated that biomimetic movement makes a robot’s movement more legible to human participants and that it is generally preferred over e.g. trapezoidal trajectories [27, 37, 56]. In human-human handovers, we know that visual feedback during the passer transport phase is crucial to enable predictive mechanisms in the physical handover [11]. A recent study showed that receiver anticipatory grip force rates scale with object mass, when the passer exhibits slightly different movement patterns at the hand level during transport of visually identical objects of different mass [42]. However, we do not know whether such effects fully transfer to robot-human interactions. First, humans are known to behave differently when collaborating with robots compared with humans [8, 40, 53]. Second, it is unclear if isolated movement patterns at the hand level are the only non-verbal cue humans use to make a prediction about the object’s mass. And, most importantly, if that information alone is sufficient in a robot-human interaction interaction.

### 2.2 Human-Robot and Robot-Human Handover

While human-human teams tend to successfully and efficiently complete object handovers, the extension to robot-human teams has proved to be challenging [52]. One reason for this lack of transferability is the bidirectional nature of interaction between two collaborating agents, requiring a thorough understanding of the human motor control system [26]. Consequently, a biomimetic approach should be most successful in designing a robot to interact with humans. The investigation of robotic movement patterns thus far has been constrained to communicating intent to hand over an object (e.g. [6]). Other research directions have included the use of speech [21] and gaze [2, 25, 47] for the same purpose.

In their recent work, Penzotti and Controzzi [54] have demonstrated a bio-inspired control law for object release, when tasking human participants to move in synchrony with the robotic passer. However, a setting of expectations of the human receiver through robotic movement was not investigated. Addressing this gap, here we aim to explore robotic motion legibility and its effect on human predictive motor control, using biomimetic cues natural to human-human interaction.

## 3 Research Goals

To enable a smooth and safe collaboration between robots and humans, it is essential that physical interactions are predictable and intuitive to humans. One promising strategy to achieve this goal is to enhance robots’ legibility by drawing inspiration from the cues humans use in their interactions. In this work, we are particularly interested in kinematic cues and their influence on human receivers’ prediction of object properties and anticipatory strategy towards a safe and effective handover.

While previous work has shown that kinematic cues can support prediction during human–human handovers, existing studies have struggled to isolate these cues from other aspects of the interaction. Indeed, no prior work has considered encoding and isolating such cues in controlled environments, none have explored simulation or robotic systems, and none have evaluated whether humans can adapt to them over time, or the influence of such adaptation between virtual and physical settings.

To address these gaps, we design an experimental framework in which the robot’s motion is fully controlled and consistent, with movement kinematics serving as the sole source of information about the object’s weight. This allows us to evaluate human predictive mechanisms in a repeatable and objective manner—measuring both behavioral adaptation and cross-platform generalization. As a result, to structure our investigation, we formulate the following research questions (RQ).

**RQ 1:** Can we design a handover task to isolate humans’ prediction about the object’s mass?

**RQ 2:** Can human participants use kinematic cues to predict an object’s properties and adapt their motor behaviour over time in response to such predictions?

**RQ 3:** Do such predictive mechanisms generalize from a virtual to a physical robot interaction?

These goals are aimed not only at understanding sensorimotor prediction in human-robot interaction, but also at offering a method for encoding communicative intent in robot motion—toward more seamless and interpretable physical collaboration.

## 4 Methods

To address our research questions, we designed controlled robot-human object handover experiments isolating kinematic information as the sole informative cue. The experimental setup was implemented in two environments, a virtual reality grasping setup [32] and a real-world physical robot, based on a service humanoid [58]. Within each environment, the object being handed over was visually identical across trials, and only the motion profile of the passer agent/robot varied with object mass.

### Task Overview

We designed handover tasks where participants acted as receivers. In both a virtual and physical settings, an agent/robot handed visually identical objects with three different weights. Only the robot kinematics varied with the object weight, while all other cues—including object appearance and release behavior—were kept constant. Participants were instructed to lift the object smoothly upon contact. Each trial involved the agent/robot lifting the object, transporting it with a mass-specific motion profile, and presenting it for grasp. We recorded grip forces and motion data to assess whether participants adapted their behavior based on the agent’s/robot’s movement alone.

### Participants

A total of 30 right-handed [51] volunteers participated in the experiments, one volunteer had to be excluded after the experiment, as they did not disclose a neurological disorder when prompted before starting. All included individuals had normal or corrected to normal vision, and to be free of acute upper limb injuries and neurological disorders. Of the remaining 29 participants, 14 took part in Virtual Agent Experiment (6 women, 8 men) aged 25 ± 3 years and 15 took part in Robotic Agent Experiment (9 women, 6 men) aged 28 ± 5 years. Before the experiment, participants provided written informed consent. This study was approved by the institutional ethics committee of the Technical University of Munich.

### 4.1 Experiment Virtual Agent

#### 4.1.1 Objective

To target RQ1 and RQ2, we isolated the kinematic movement of a simulated passing agent as a cue and measure participants’ anticipatory grip forces as a metric of predictive motor behaviour.

#### 4.1.2 Experimental Setup

In this experiment, we devised a setup with two haptic robots (Phantom Premium 1.5 HF, 3D Systems, Rock Hill, USA), providing position and force feedback, and a monitor-mirror system for visual feedback integrated to a virtual environment. This setup was similar to our previous work [32]. The two haptic robots were connected to the participant’s index finger and thumb respectively using a 3D printed thimble and rigid medical tape. The view of their hand was blocked by the mirror (see Fig. 2A). Participants were seated in a chair in front of the system in a fixed position. In the experiment, participants viewed a virtual environment based on CHAI3D [10] and Open Dynamics Engine libraries [57], validated for human object manipulation experiments [32]. The participant’s finger tips were represented by gray spheres with a radius of 1cm (Fig. 2B). Within the environment, positions and forces were sampled at 500Hz, visual feedback was provided at 60Hz.

**Fig. 1.**
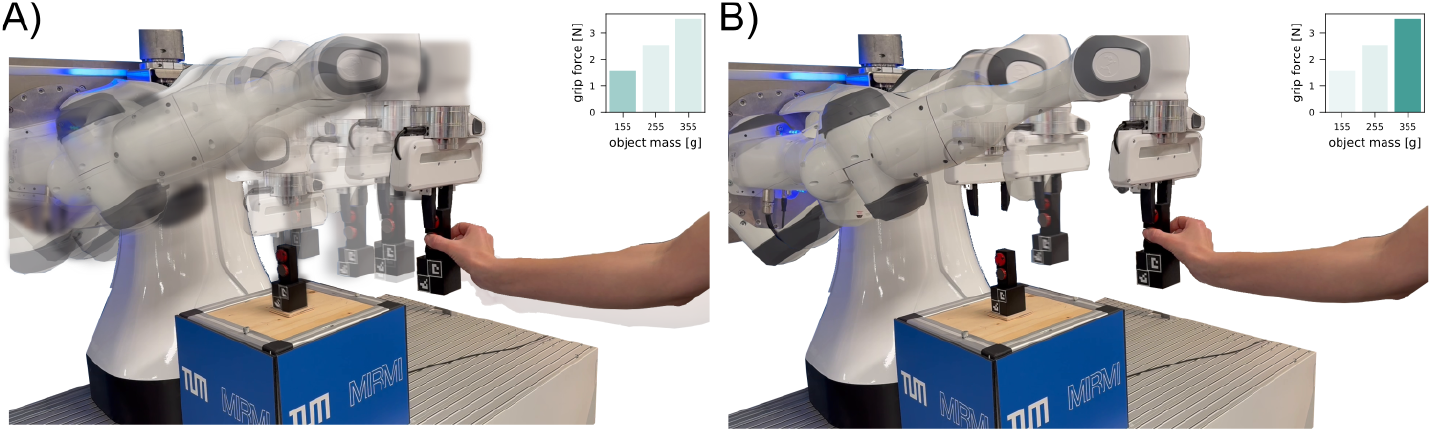
We use biomimetic robotic kinematic cues to enable human predictive control in a robot-human object handover. A) Fast robotic movement leads to low anticipatory grip forces. B) Slow robotic movement leads to high anticipatory grip forces.

**Fig. 2.**
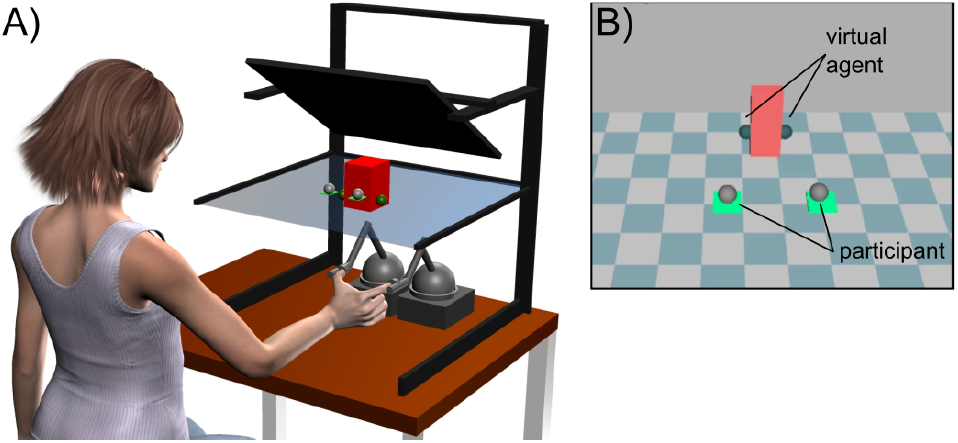
Experimental setup for Experiments with virtual agent. A) Virtual grasping setup used in Virtual Agent Experiment. Participants’ index and thumb were connected with medical tape to a haptic robot each to provide position and force feedback. B) View of the screen providing visual feedback of the virtual scene including cursors of the simulated agent and the participant.

In the environment, a rigid (stiffness=700N/m), box-shaped test object (2×4×10cm) with uniform mass distribution was used. Depending on the condition, the object weight varied from 100, 200, 250, and 300g.

#### 4.1.3 Experimental Paradigm

A virtual agent represented by two teal-colored cursors was the passer, while the participant was the receiver. Each trial started with the participant placing their index finger and thumb on two starting platforms, after 500ms, the agent’s cursors appeared and reached toward the object, grasped, held and lifted it up by 10cm and transported it toward the participant by 30cm. During this action, the participant was required to remain in the starting positions, if they failed to do this the trial was reset and restarted. Once the handover position between the participant’s fingers was reached, the agent and object remained stationary and locked in space for 2s during which participants could grasp and hold the object. After this time, the cursors released the object and gravitational acceleration acting on it was linearly ramped up over 100ms. After another 2s, a plane appeared 3cm above the handover position. Participants were asked to lift the object above the plane, release it, and return to the start positions for the next trial.

Each of the agent’s point-to-point movements (reach, lift, and transport) was modeled according to the minimum jerk model, which approximates human point-to-point movement [20],

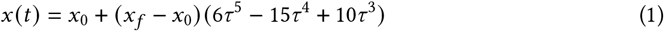

where *x*_0_ and *x*_*f*_ are the initial and final position, respectively, and *τ* is the normalized time, given by 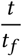, where *t_f_* is the final time. The duration for each phase was adapted from previous work [13] and modulated with the object weight. That is, a heavy object was matched with long durations and a light object matched with short durations (see Table 1). The mass-trajectory pairings remained consistent across the experiment.

**Table 1.** Duration of agent movement phases in virtual agent experiment for different trial phases, adapted from [13]. The duration for each phase where the agent is interacting with the object is increased with increasing object mass. The gray column indicates practice configuration.

| Phase | Duration in s |  |  |  |
| --- | --- | --- | --- | --- |
| Object Mass in g | 100 | 200 | 300 | 250 |
| Reach | 1.1 | 1.1 | 1.1 | 1.1 |
| Grasp & Hold | 0.0176 | 0.176 | 0.8800 | 0.4400 |
| Lift | 0.4380 | 0.8760 | 1.3140 | 1.0950 |
| Transport | 0.5020 | 1.0040 | 1.5060 | 1.2550 |

#### 4.1.4 Experimental Procedure

Participants completed four experimental parts: Practice, Random 1, Blocked, and Random 2 (Fig. 3D). Before starting the practice phase, participants familiarized themselves with the virtual setup by lifting and replacing an object ten times using their right hand and our virtual object manipulation setup [32]. In the practice part of the experiment, participants completed two blocks of 25 trials each. In this part, the test object’s mass and the agent movement trajectory were consistent. The Random 1 phase consisted of six blocks with 18 trials each. Within each block of 18 repetitions, each of the three possible mass-trajectory combinations was presented six times in a pseudo-random order. Therefore, in a set of three trials each mass-trajectory combination was presented once. The Blocked phase consisted of three blocks of 18 trials where the mass-trajectory combination was constant for all 18 consecutive trials of a block. Finally, the Random 2 phase again consisted of three blocks with 18 trials each (randomized as in Random 1 phase). Between participants, the order of mass-trajectory blocks in the Blocked part of the experiment was pseudo-randomized and counter-balanced across participants. For the rest of the experiment, the order of trials was identical for all participants. At the end of the full experiment, participants were asked to fill out a questionnaire, that included the following questions:

**Fig. 3.**
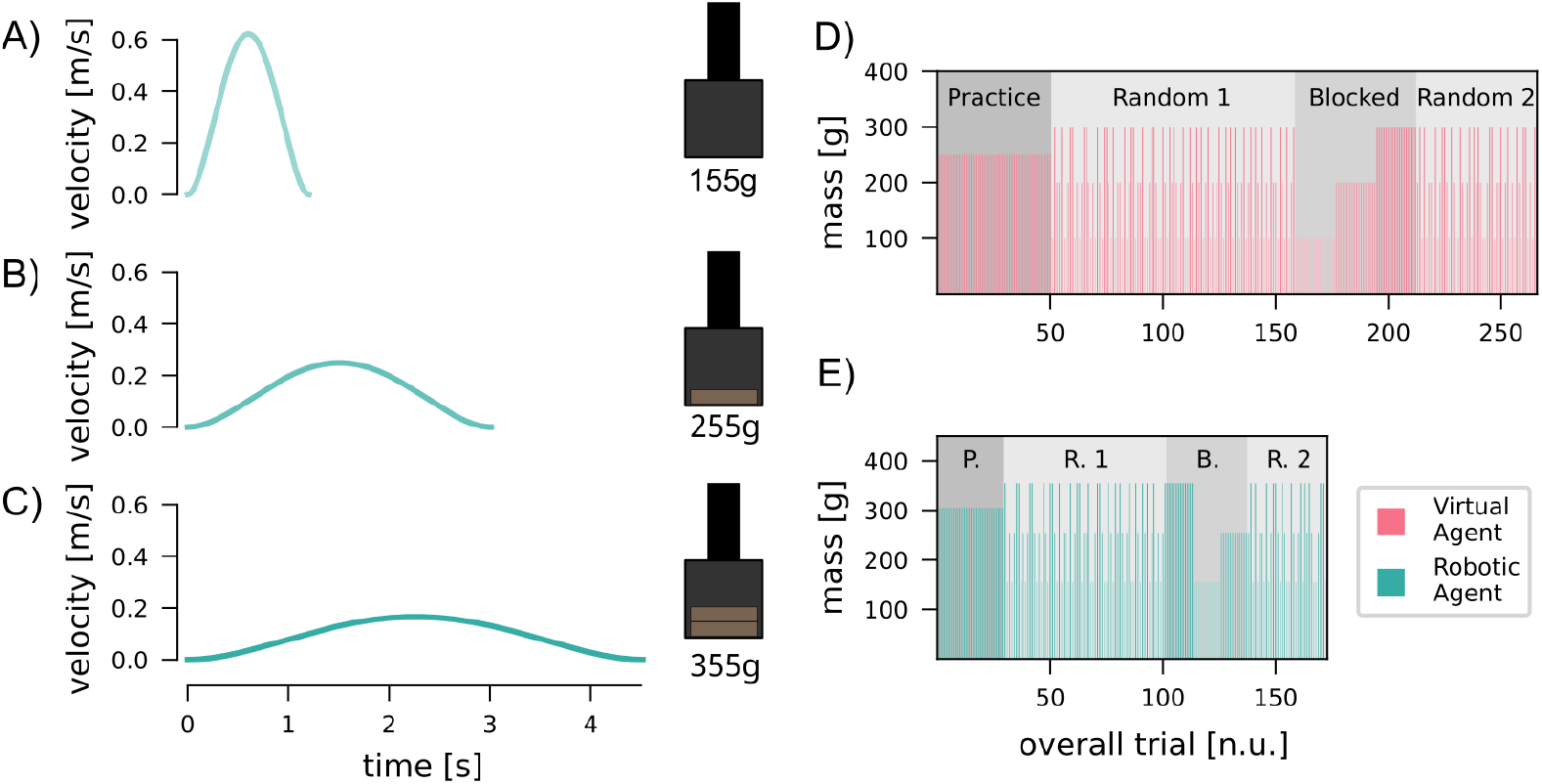
Methods for both experiments. A)-C)) Velocity profiles over time for the transport phase of the three object weights presented in the robotic agent experiment. A higher peak velocity is matched with a lower object weight; a lower peak velocity is matched with a higher object weight. D) Experimental Design in Virtual Agent Experiment. Task practice was followed by an initial random part, a blocked part, and a second random part. The height of the bars indicates object weight in individual trials. E) Experimental Design for Robotic Agent Experiment. Experimental design is the same as Virtual Agent Experiment, with lower repetitions for each phase.

- Did the object have different weights? If yes, how many different weights across the experiment?
- Did you notice different agent behavior between trials? If yes, what was it?
- Did you notice a connection between the object weights and the agent behavior? If yes, what was it?

#### 4.1.5 Data Analysis

Participants’ grip and load forces were calculated based on the forces produced by the haptic robots and object orientation in the virtual environment. Grip force (**GF**) was defined as the normal component of digit force with respect to the object’s surface. Load force (**LF**) was defined as the vertical component of the digit force, normal to the ground. Both forces were filtered with a 6th-order zero-phase butterworth filter with 20Hz cutoff. Finally, we recorded the vertical object position throughout the trial. Using these data, we detected the following metrics for each trial.

- **GF**_*ant*._ – Anticipatory Grip Force;
- **GF**_*static*._ – Static Grip Force;
- **GF**_*corr*._ – Grip Force Correction;
- **LF**_*ant*._ – Anticipatory Load Force;
- **disp**_*vert*.,*max*_ – Maximal Vertical Object Displacement.

We define **GF**_*ant*._ as the mean grip force in the last 20 ms before the start of the release of the test object and **GF**_*static*_ as the mean grip force 750 ms after the release of the object to 500 ms before the end of the trial. The variable **GF**_*corr*._ is the difference between *GF*_*ant*._ and *GF*_*static*._. A positive value indicates an increase in grip force from the predictive to the static phase. Finally, **LF**_*ant*._ is the mean load force in the last 20 ms before the start of the release of the test object, and **disp**_*vert*.,*max*_ is the maximum vertical deviation of object position from handover position after the passer has released it.

To intuitively summarize values across the three presented objects in the Random 1 and 2 phases of the experiment, fit a linear regression, using

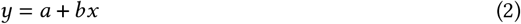

where *a* and *b* are the intercept and slope, respectively. The x-values were calculated by

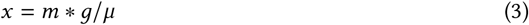

where *m* is the object mass in kg, *g* is the gravitational acceleration constant 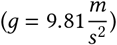, and *μ* was the friction between object and finger (here *μ* = 1). To remove trial-by-trial noise, we smoothed the data for each object mass using a moving average filter with size 6 before fitting a regression to each trial. The parameters of this linear regression indicate how well participants can predictably scale the anticipatory grip force to the upcoming object, with higher slope and lower intercept indicating better adaptation to the respective object mass.

To determine the trial in phase Random 1 at which participants no longer adapted to a specific cue, i.e. where they had reached a steady-state performance, we used the expanding window protocol as introduced in [1]. Namely, we used the previously calculated intercept values for the Random 1 phase of the experiment and raw data for the Practice phase of the experiment as an input. We then iteratively fit a linear regression to this data using an expanding window approach, starting at the last trial and with a window of 15 trials. For each fit, we recorded the confidence interval. The first steady-state trial was defined as the trial corresponding to the lowest recorded confidence interval across the full phase.

Some of the data presented for this experiment have previously been reported in [30].

### 4.2 Experiment Robotic Agent

#### 4.2.1 Objective

Here, we tested whether our results from the VR experiment would translate to a handover with a physical robot. More specifically, we address RQ3 to investigate whether the adaptation of grip forces to the robot kinematic cue is affected by interacting with a physical robot and object in the real world.

#### 4.2.2 Experimental Setup

In this experiment, the setup consisted of the upper-body bimanual-manipulation system of the service robot GARMI [58], an instrumented test object, an RGB camera, and three PCs, one for robot control, one for streaming grip force sensor data, and one for data saving (Fig. 4). The bimanual robot consisted of a two vertically-mounted 7 degrees of freedom Franka robot arms with modified gravity compensation force (due to the change in dynamics), equipped with the standard Franka gripper [33]. During the experiment, only the right arm was used. The left arm was in a neutral, stationary position throughout the experiment. The pose was chosen according to the most preferred pose by users found in previous work [48]. We recorded the right arm’s position, torque, and end-effector force throughout at 100 Hz (streamed to recording PC via ROS).

**Fig. 4.**
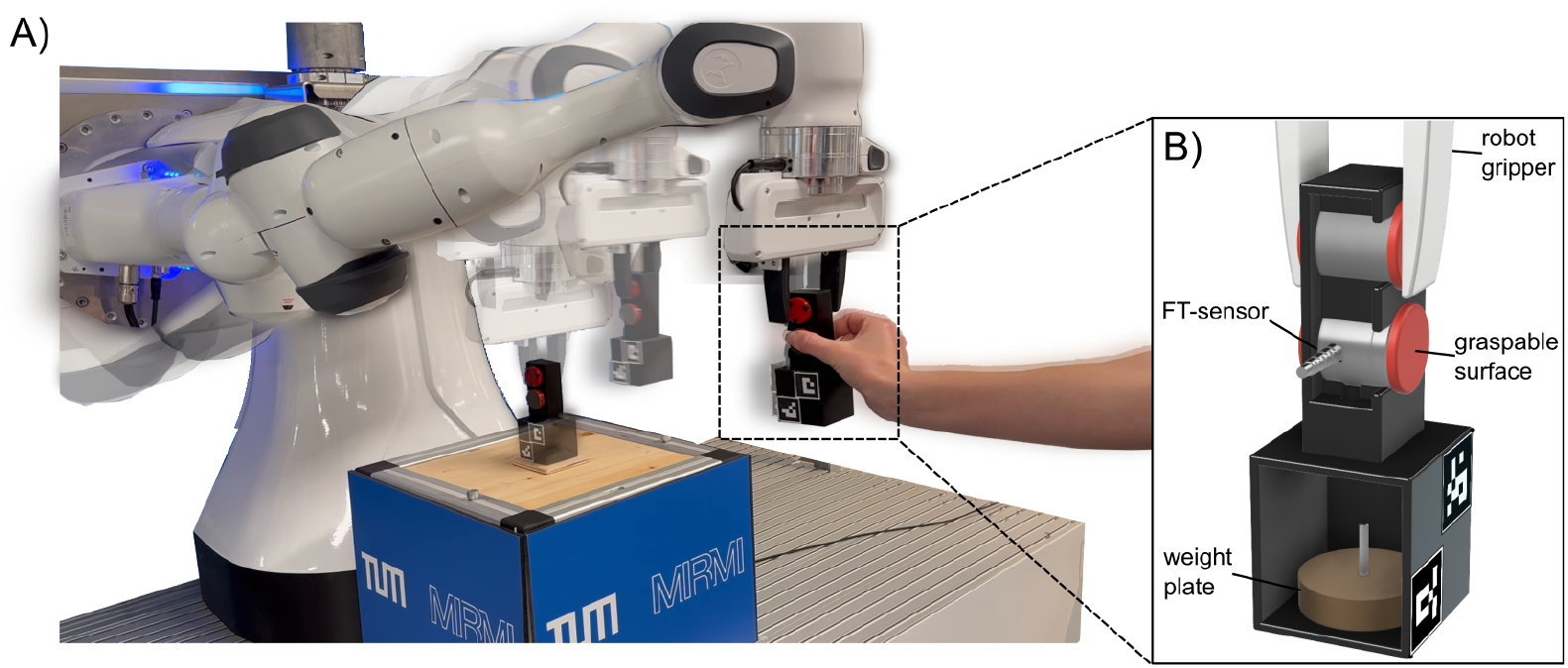
Experimental setup for Experiments with robotic agent. A) Experimental setup in Robotic Agent Experiment. Participants sat in front of a reduced version of the service robot GARMI. GARMI handed a test object at a fixed position to the participant. B) The test object held by the robotic gripper. A force sensor with graspable surface for the participant is connected to the upper part of the object. The bottom part of the object includes a slot to insert weight plates and had two ARUCO markers attached to the side visible to a camera.

The test object (Fig. 4B) was a 3D printed plastic structure consisting of two box-shaped parts. The upper part (outer dimensions 79x25x35mm) included the grasp surface for the robot gripper and a slot for a six-axis force/torque sensor (ATI Nano25-E). P800 sandpaper was applied to the grasp surfaces to standardize the object friction between participants. Sensor data was recorded at 250Hz and streamed to the recording PC via ROS. The bottom part (outer dimensions 56x51x51mm) included a slot in the back, that allowed the experimenter to insert weights to change the weight of the object, without changing its appearance. The two parts of the object were connected with screws to form one rigid object.

Further, we used an RGB camera (Logitech C922 Stream Pro) to track the object position after it was released by the robot. The camera was running at 30fps and images were streamed to the recording PC via a ROS 1-based message passing system [55]. Two ARUCO markers were attached to the side of the part facing a camera to track object position.

Participants sat on a chair in a fixed position in front of GARMI. GARMI’s movement was implemented using a polynomial time scaling law, (1), as in [44]. The end-effector orientation was kept fixed throughout the motion generation process. The end-effector reached the end of the table GARMI was mounted on while still respecting the robot workspace limits. For the lower-level tracking controller, a proportional feedback control law (with a gain value of 10) was implemented at the velocity level. As far as stability is concerned, this control strategy achieves exponential convergence to the desired reference pose [17].

#### 4.2.3 Experimental Paradigm

In this object handover experiment, GARMI was the passer, while the participant was the receiver. Each trial was manually started by one of the experimenters by pressing a button. From the reset position, GARMI’s right arm reached toward the object, grasped, held, and lifted it up by 13cm, and then toward the participant by 40cm. Consequently, the movement consisted of phases for reach, grasp and hold, lift, and transport. Once the handover position was reached, the robot arm and hand remained stationary for 2s before the gripper was opened. Participants were asked to grasp and hold the test object using their right index finger and thumb, before the gripper opened. After 2 more seconds, the robot arm retracted and moved towards its reset position. Once the retraction movement started, participants were asked to pass the object to the second experimenter, who changed the object’s weight for the next trial and replaced the object on the table. The participants were aware of the weight change, but were unable to see the weight for the next trial.

Analogous to the Virtual Agent Experiment, each of the robot’s point-to-point movements (reach, lift, and transport) was modeled according to the minimum jerk model. The duration for each phase was adapted from previous work [13, 30]. The phase duration was modulated with the object weight (Fig. 3 A-C), with a heavy object matched with long durations and a light object matched with short durations (see Table 2). The mass-trajectory pairings remained consistent across the experiment.

**Table 2.** Duration of robotic movement phases in Robotic Agent Experiment for different trial phases, adapted from [13]. The duration for each phase where the agent is interacting with the object is increased with increasing object mass. The gray column indicates practice configuration.

| Phase | Duration in s |  |  |  |
| --- | --- | --- | --- | --- |
| Object Mass in g | 155 | 255 | 355 | 305 |
| Reach | 2 | 2 | 2 | 2 |
| Grasp & Hold | 1.047 | 1.468 | 3.340 | 2.170 |
| Lift | 1.314 | 2.628 | 3.942 | 3.285 |
| Transport | 1.506 | 3.012 | 4.518 | 3.765 |

#### 4.2.4 Experimental Procedure

Participants completed the same experimental phases as in the Virtual Agent Experiment: Practice, Random 1, Blocked, and Random 2 (Fig. 3 H). Before starting the practice phase, participants familiarized themselves with the test object by lifting and replacing it on a table ten times using their right hand and by lifting it using their right hand and letting it slip into their left hand ten times. In contrast to Virtual Agent Experiment, we reduced the number of trials by one third, as our previous work showed that motor adaptation in object manipulation occurs slower in VR compared to the physical world [32]. Consequently, the practice part of the experiment, consisted of two blocks of 15 trials each. The Random 1 phase consisted of four blocks with 18 trials each. The Blocked phase then consisted of three blocks of twelve trials where the mass-trajectory combination was constant for all twelve consecutive trials of a block. Finally, the Random 2 phase consisted of two blocks with 18 trials each, where the block order was again random (same as in Random 1 phase).

Between participants, the order of mass-trajectory blocks in the Blocked part of the experiment and the initial three trials in Random 1 was pseudo-randomized and counter-balanced across participants. For the rest of the experiment, the order of trials was identical for all participants. Importantly, throughout the entire experiment, regardless of whether the weight was changed, the second experimenter “changed the weight”, such that this did not provide a specific cue about the upcoming trial. At the end of the full experiment, participants were asked to fill out a questionnaire, that included the following questions:

- Did the object have different weights? If yes, how many different weights across the experiment?
- Did you notice different robot behavior between trials? If yes, what was it?
- Did you notice a connection between the object weights and the robot behavior? If yes, what was it?

#### 4.2.5 Data Analysis

Data processing, data analysis and statistics were performed offline in Python. Grip forces, as measured by the force-torque sensors, were filtered with a 15Hz lowpass filter (zero-phase 8th order butterworth). End-effector forces of the robot arm were low pass filtered at 10Hz (zero-phase 6th order butterworth). Finally, the object position was recorded using a RGB-camera and two ARUCO markers on the object. For each frame, we detected the two markers, offset corrected their positions, and applied an IQR (interquartile range) filter with a 10 sample window replacing values larger than 1.5 × *iqr*_*window*_ with interpolation to remove outliers.

Using these data, we detected the following metrics for each trial:

- ***GF*** _*ant*._ – Anticipatory Grip Force;
- ***GF*** _*static*._ – Static Grip Force;
- ***GF*** _*corr*._ – Grip Force Correction;
- ***LF*** _*ant*._ – Anticipatory Load Force;
- ***disp***_*vert*.,*max*_ – Maximal Vertical Object Displacement.

We define ***GF*** _*ant*._ as the mean grip force in the last 20ms prior to the opening of the gripper and ***GF*** _*static*._ as the mean grip force 750ms after the opening of the gripper to 500ms before the end of the trial. The ***GF*** _*corr*._ is the difference between ***GF*** _*ant*._ and ***GF*** _*static*._, while the ***LF*** _*ant*._ is computed by

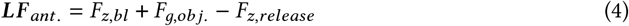

where *F*_*z,bl*_ is the baseline end-effector force as measured by the robot in the z-axis (without load), *F*_*g,obj*._ is the gravitational force of the test object, and *F*_*z,release*_ is the mean force in the z-axis in the last 20ms prior to the opening of the gripper. Finally, after the gripper opened, and before the robot arm retracted, i.e. the object was handed back to the experimenter, we measured the maximum vertical deviation of the object ***disp***_*vert*.,*max*_.

Additionally, using data acquired in the object familiarization, we detected the slip force, that is, the minimum grip force applied before the object slipped from participants’ grasp. Using the slip force, we estimated the friction between the object and finger.

Similar to the Virtual Agent Experiment and as described above, we fit a linear regression to the outcome metrics to summarize values across the three presented object masses. To determine the first steady-state trial, we used the method described in [1] with an initial window size of 10 samples.

### 4.3 Statistical Analysis

To test the effect of movement velocity on memory formation and the performance metrics within each experiment, we used a repeated-measures ANOVA with “object mass” (3 levels) as the independent factor. In the case of non-sphericity, we corrected the RM-ANOVA using the Greenhouse-Geisser correction. If the main effects of the RM-ANOVA were statistically significant, the specific differences were compared using a paired t-test with Bonferroni correction. Statistical significance was determined at the p<0.05 threshold in all tests.

## 5 Results

### 5.1 Task Proof of Concept

In both experiments, participants were repeatedly handed an object with varying mass. To validate the general task design, we evaluate the questionnaire participants answered after completion of the experiment. Further, to demonstrate the validity of our task to measure human predictions, we analyzed the trial-by-trial adaptation in the experimental phases where the object mass was constant for multiple repetitions (Practice and Blocked).

#### 5.1.1 Virtual Agent Experiment

In this experiment, participants were repeatedly handed an object with a different mass by an artificial agent in a custom virtual environment. All participants successfully completed the task, with 420 trials or 1.6% failed trials across all participants where the object was dropped. Out of 14 participants, all noticed the different object weight and agent trajectories. 12 participants reported having noticed the connection between weight and behaviour, i.e. the movement velocity, only these participants were included in the further analysis.

After a few initial trials in the Practice phase, participants were able to hold the object without dropping it and iteratively reduced their anticipatory grip forces (Fig.5A). This adaptation process lasted on average 11 repetitions. Within the Blocked phase of the experiment, participants readily adapted their motor behaviour to the object weight. Their anticipatory grip force was scaled according to weight with significantly lower anticipatory grip forces for the lightest mass compared to the medium and heavy objects (Fig. 5B, Table 3). Similarly, the static grip force showed clear scaling to object mass with all comparisons between masses showing significant differences (Table 3). When subtracting the predictive component from the static grip force, we observe an upscaling of grip force, that is, we find that all values for corrective grip force are positive (Fig. 5C). Therefore, rather than decreasing their grip force when the object is released and they receive feedback on the object mass, they consistently applied higher forces (3).

**Table 3.** Summary of RM ANOVA and bonferroni corrected post-hoc paired t-tests of the virtual agent experiment. Significant comparisons are marked in bold text.

|  | RM ANOVA | Post-hoc T-test |  |  |
| --- | --- | --- | --- | --- |
|  | Main Effect | 100-200g | 100g-300g | 200g-300g |
| Blocked |  |  |  |  |
| $GF_{ant.}$ | $F_{(1.07,11.73)} = 15.95, p = 0.002$ | **p<0.001** | **p=0.003** | p=0.052 |
| $GF_{static}$ | $F_{(1.10,12.10)} = 26.23, p < 0.001$ | **p<0.001** | **p<0.001** | **p=0.019** |
| $LF_{ant.}$ | $F_{(1.88,20.66)} = 49.87, p < 0.001$ | **p=0.001** | **p<0.001** | **p<0.001** |
| $disp_{vert.,max}$ | $F_{(1.19,13.14)} = 37.21, p < 0.001$ | **p<0.001** | **p<0.001** | **p=0.002** |
| Random 1 - Steady-State |  |  |  |  |
| $GF_{ant.}$ | $F_{(1.07,11.75)} = 8.15, p = 0.014$ | **p=0.048** | **p=0.042** | p=0.061 |
| $GF_{static}$ | $F_{(1.11,12.24)} = 24.37, p < 0.001$ | **p<0.001** | **p<0.001** | **p=0.005** |
| $LF_{ant.}$ | $F_{(1.50,16.52)} = 1.40, p = 0.269$ | n.a. | n.a. | n.a. |
| $disp_{vert.,max}$ | $F_{(1.17,12.87)} = 50.01, p < 0.001$ | **p<0.001** | **p<0.001** | **p<0.001** |

**Fig. 5.**
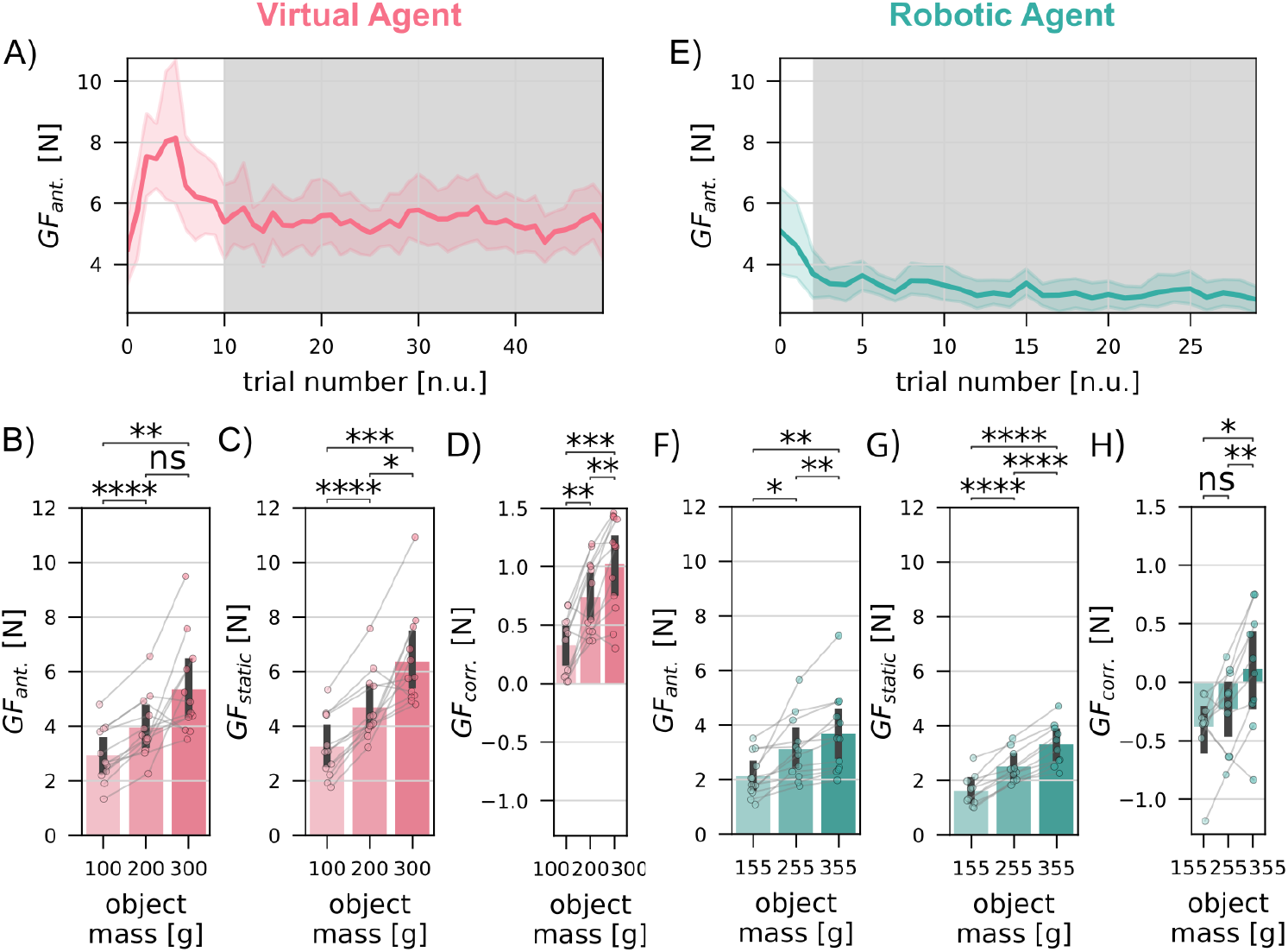
Adaptation of anticipatory grip force to consistent objects. A) & E) Mean trial-by-trial and 95% confidence interval in practice phase of Virtual Agent Experiment (A) and Robotic Agent Experiment (E). The steady-state is indicated by gray background shading. B) & F) Averaged anticipatory grip force values for different object masses in the blocked phase of the Virtual Agent Experiment (B) and Robotic Agent Experiment (F). C) & G) Averaged static grip force values for different object masses in the blocked phase of Virtual Agent Experiment (C) and Robotic Agent Experiment (G). D) & H) Averaged grip force correction values for different object masses in the blocked phase of Virtual Agent Experiment (D) and 2 (H). B)-D) and F)-H) Individual data points show participant data, bars and error bars represent the mean and 95% confidence interval.

#### 5.1.2 Robotic Agent Experiment

All human receivers successfully completed handovers from the robotic passer, which moved slow, medium and fast for heavy, medium, and light objects, respectively. Out of the 15 participants, all noticed the different object weight and robot trajectories. 12 participants reported noticing the connection between weight and robot trajectory. Only those participants who noticed the connection were included for further analysis.

Within the Practice and Blocked parts of our experiment, participants were repeatedly handed an object with the same weight and trajectory. On average, participants generated a stable anticipatory grip force after being handed the test object by the robotic agent five times (5A). Therefore, they were able to quickly adapt to the task and scale the grip force to task demands in a predictive way. When averaging across all 12 repetitions for each mass in the Blocked phase of the experiment, we observe scaling of anticipatory grip force according to the object mass (Fig.5B). Similarly, the static grip forces show strong scaling of grip force according to object mass after full haptic feedback was available. Comparing between predictive and static grip forces, we find minimal adjustments, with a slight reduction in grip force for the low and medium object masses.

All statisical comparisons between masses show significant differences for the anticipatory and static grip forces, indicating that participants clearly adapt their predictive behaviour to the task demands when the same object mass is presented repetitively (Tabs.3&4).

### 5.2 Cue-based Prediction of Different Object Masses

After an initial familiarization to the task, participants were repetitively handed objects of varying masses in a random order. In contrast to the Practice and Blocked phases participants could not use the previous trial to predict the object mass, but had to instead rely on the kinematic cue given by the virtual agent or the robotic agent, respectively. Therefore, a scaling of anticipatory grip force to the respective object mass, and the strength and quality of this scaling effect, would indicate the existence of and certainty in a memory correlating movement kinematics with object mass.

#### 5.2.1 Virtual Agent Experiment

We applied a linear regression fit to outcome metrics to summarize values for all three mass conditions, into slope and intercept. Based on these values, we ran the expanding window protocol introduced by Blustein et al. [1], to estimate adaptation and steadystate trials. On average, participants adapted their predictive grip forces to the movement cue after 18 repetitions of each object (Fig.6A&B). After successful adaptation, we observed significant differences in scaling for the light object compared to medium and heavy objects, but not between the medium and heavy objects (Fig.6E). Participants were, on average, faster to scale their static grip force to the object weight (17 repetitions), an effect that is immediate in simple, real-world object lifting tasks [59]. Similar to the Blocked experimental phase, we observe an increase in grip force values (positive *GF*_*corr*._) after release of the object by the agent (Fig.6I). This correction significantly scaled with the respective object mass (Tab. 3).

While the average slope is slightly higher in the Blocked compared to Random experimental phases (Fig.6A), there is no visible change between the random phases. Therefore, the adaptation to the cue-weight pair was saturated and no deadaptation was induced by the Blocked phase. This pattern is consistent with the intercept values that are slightly lower in Blocked, but consistent between end of Random 1 and Random 2.

**Fig. 6.**
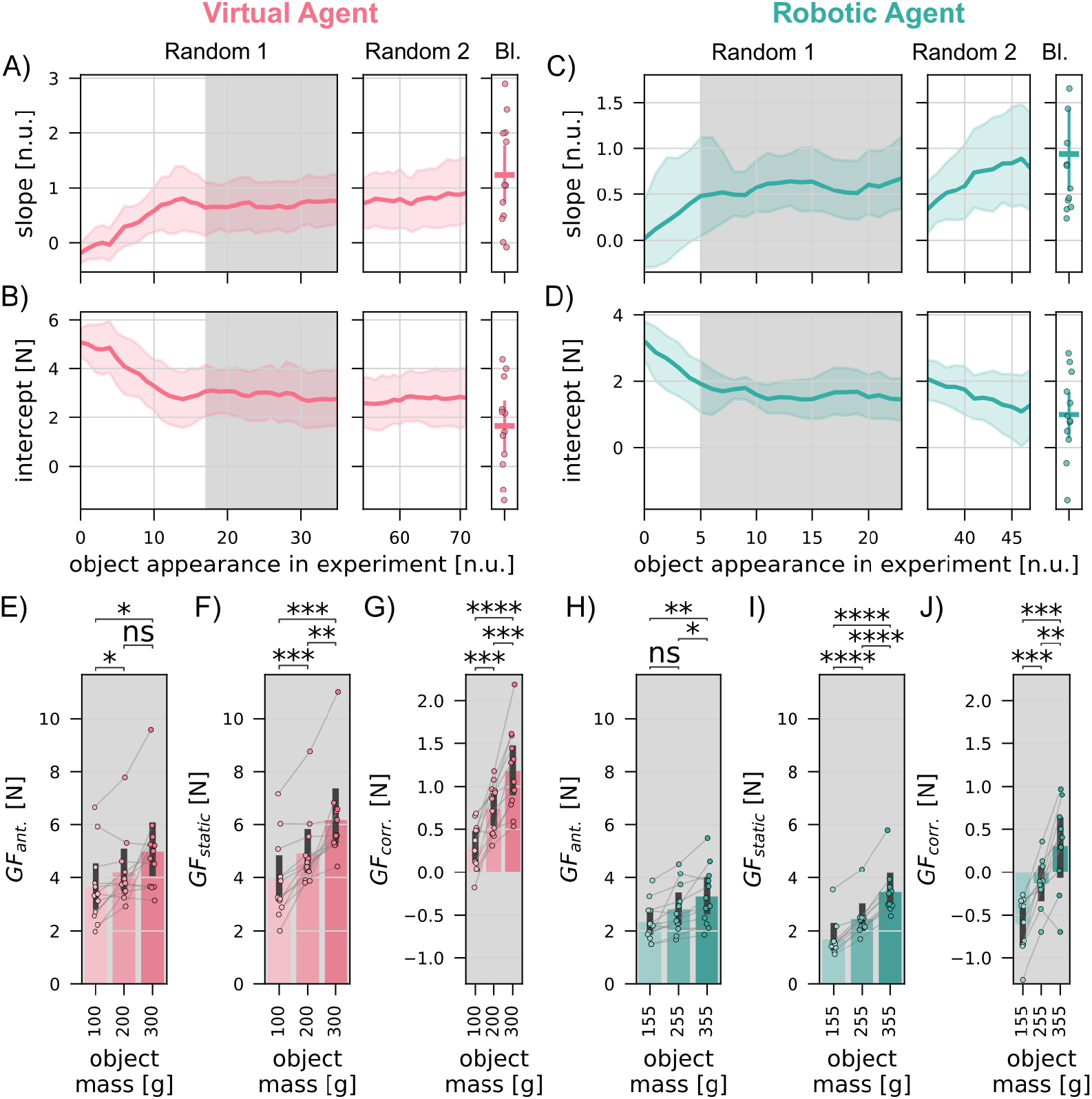
Smoothed trial-by-trial data for all experimental phases in Virtual and Robotic Agent Experiments. A&B) show smoothed trial-by-trial mean and 95% confidence interval for slope and intercept values across all participants for Virtual Agent Experiment. C&D) show smoothed trial-by-trial mean and 95% confidence interval for slope and intercept values across all participants for Robotic Agent Experiment. The shaded areas indicate the steady-state trials in Random 1 phases. For reference, we also show the results for the Blocked phase of the experiment (Bl.). One participant’s slope in the Blocked phase is off scale and was cut from the figure to preserve the overall scaling. The participant’s data is included in all mean and ci values. E-J) Averaged anticipatory grip force, static grip force and grip force correction values for different object masses in steady-state trials of the random phase of Virtual Agent Experiment (E-G) and Robotic Agent Experiment (H-J). Individual data points show participant data, bars and error bars represent the mean and 95% confidence interval.

#### 5.2.2 Robotic Agent Experiment

On average, participants adapted their anticipatory grip forces for the initial six trials, and remained stable for the remaining repetitions (Fig.6A), meaning that, each object mass was presented six times before a stable prediction was reached. Within the steady-state repetitions, the anticipatory grip force shows clear scaling to object mass (Fig.6C & Table4, Random 1) between the light and heavy weight and the light and medium weight, but not between light and medium weights. Similarly, participants adapted their static grip forces to the object mass for six repetitions per object mass. After this adaptation, we observe clear scaling to the respective object mass (Fig.6I). Similar to the Blocked phase, the correction of grip force, that is, the difference between static and anticipatory grip force is generally low. On average, we observe a slight reduction for the lowest object mass and a slight increase for the highest object mass (Fig.6J).

The trial-by-trial average slope shows little change between the end of Random 1 and the beginning of Random 2, however, as observed in Virtual Agent Experiment, the slope in the Blocked phase of the experiment is higher than in both random phases. Similarly, the intercept shows little variation between the random phases, but is decreased in the blocked phase.

**Table 4.**
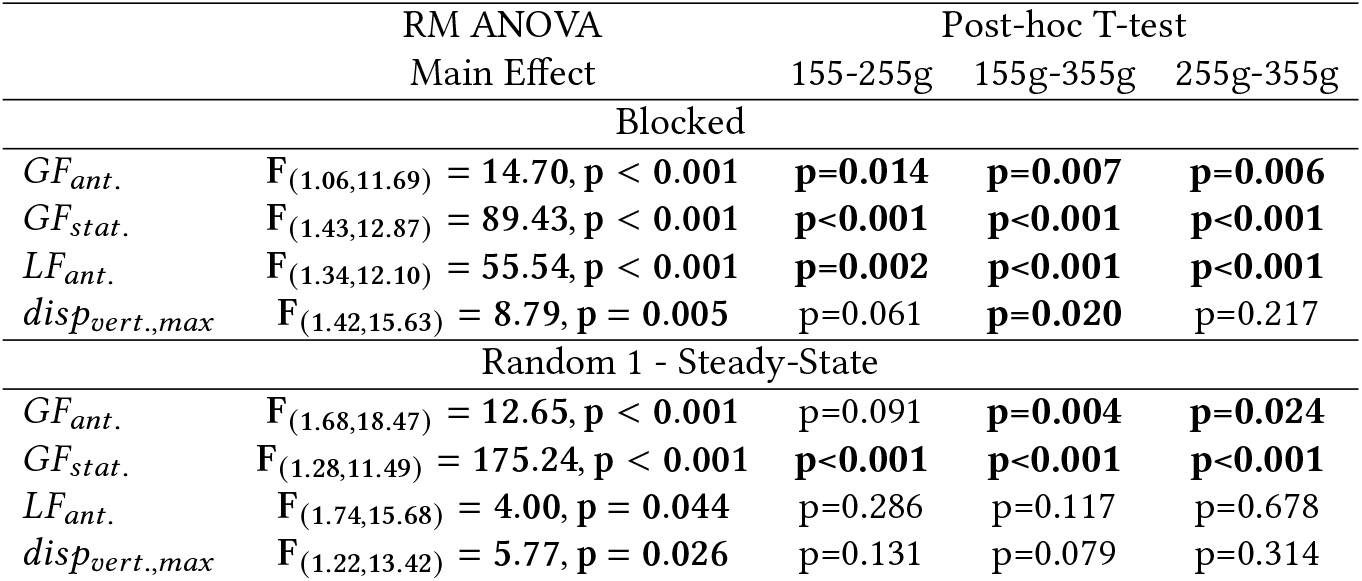
Summary of RM ANOVA and bonferroni corrected post-hoc paired t-tests of robotic agent experiment. Significant comparisons are marked in bold text.

### 5.3 Quantitative Comparison between Experiments

When comparing outcome metrics of both experiments, we observe an overall similarity in adaptation patterns. In the blocked phase of the experiment, participants reliably scale the grip forces to the task demands. That is, we observe significant differences between forces applied for each object mass. In both experimental phases, participants adapted faster in the real-world experiment in contrast to the virtual reality. Interestingly, while the slopes for the anticipatory grip force in both experiments converged to very similar values (Fig.7A), the intercept values in the virtual agent experiment remained elevated even after convergence (Fig.7E). To better understand the adaptation process and differences between Experiments 1 and 2, we fit exponential decay functions to the slope and intercept values for anticipatory grip force. For the both metrics (slope and intercept) we employed a leave-two-out bootstrapping method on the existing dataset to generate additional data points for our model. The slope data was fit to

**Fig. 7.**
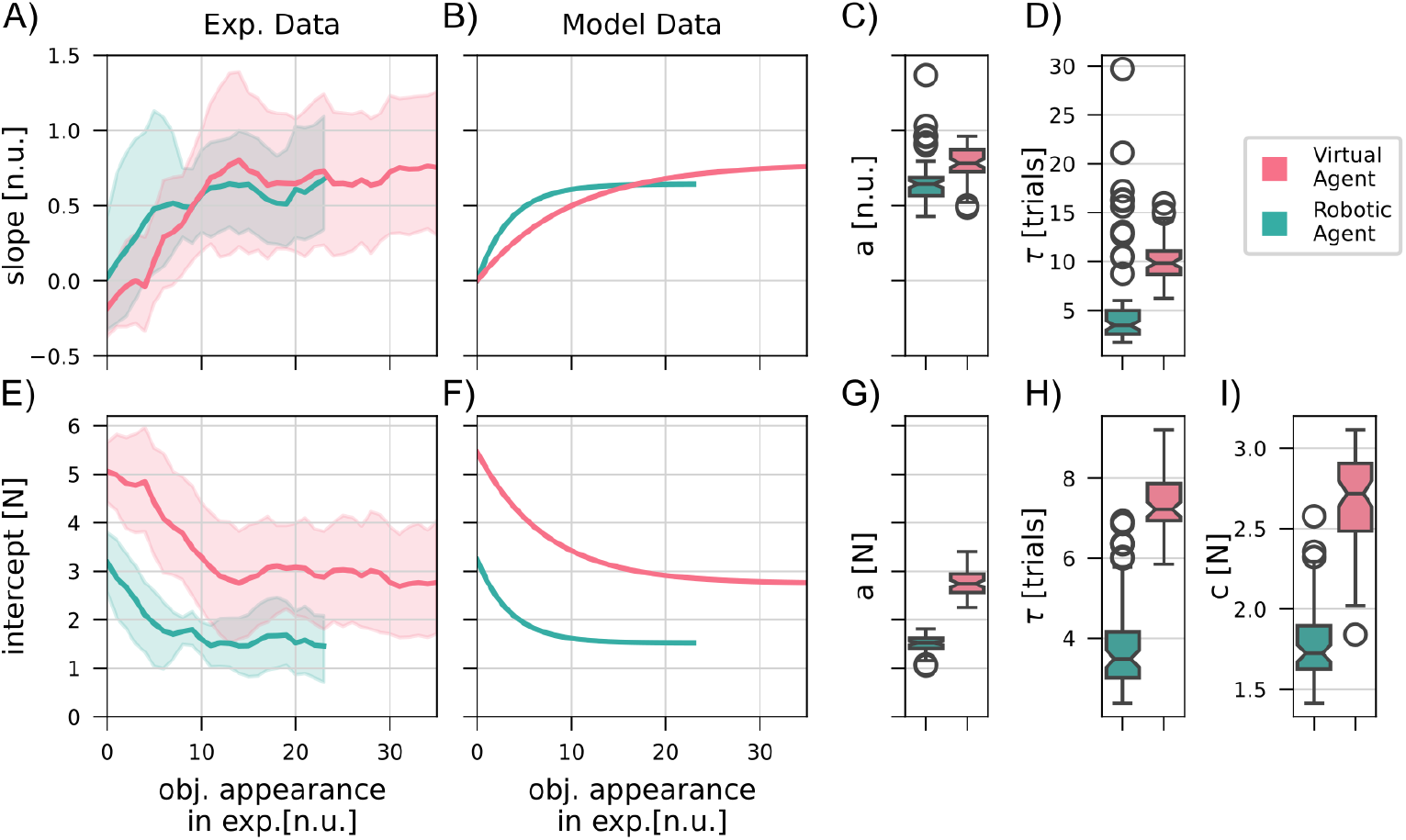
Smoothed trial-by-trial adaptation across participants in random 1 and 2 and model results. A) and E) show slope and intercept values for anticipatory grip force in the two experiments. B) and F) show best fit exponential functions to the median metric values. C-D)&G-I) show median and iqr for fitted model parameters based on bootstrapped data.

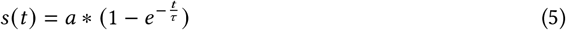

where *a* is the asymptote (i.e. the final slope the function approaches), and *τ* is the number of trials to reach full adaptation. The intercept data was fit to

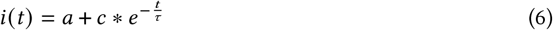

where *a* is the asymptote (i.e. the final slope the function approaches), *c* is the expected change in intercept during training and *τ* is the number of trials to reach full adaptation. Using these functions, the anticipatory grip force at a trial can be described as

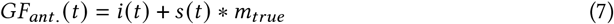

where *m*_*true*_ is the current object mass.

Our model results (Fig.7) highlight the increased adaptation process in virtual reality compared to the real world. This is reflected in increased values of *τ* for the virtual agent experiment in both metrics; slope and intercept. While the asymptote *a* is similar in the adjusted slope, we observe an offset of 1.22N for the virtual agent intercept value. Finally, while participants in virtual reality converged to a higher intercept value, they also demonstrated a higher change in intercept value from the initial to the adapted trials. Here, we observe a median value of 2.72N for the virtual agent experiment in contrast to 1.72N for the robotic agent experiment.

### 5.4 Additional Behavioural Metrics

Additional to the grip force, which is our primary metric of prediction, we measured the vertical object deviation after the passer release and the anticipatory load force to gain a better understanding of how human participants adapt motor commands in our specific task.

#### 5.4.1 Maximum Object Deviation after Passer Release

In general, participants in both experiments were able to quickly reduce the amount of vertical object displacement after the handover in the random phases of the experiment. In both experiments, participants required, on average, six repetitions per object mass to reach a steady-state. Within this steady-state, the slope for the virtual agent experiment converged to a value of -1, whereas the slope for the robotic agent experiment converged to a value of 0 (Fig.8A). The intercept values, converged to 1cm and -1cm for the virtual agent and robotic agent experiments, respectively (Fig.8B). Therefore, in both experiments, the movement after the handover action was minimal. These patterns were consistent with that observed in the Blocked experimental phase. Despite the low values, the repeated measures ANOVA revealed significant comparisons in both Experiments in both the Blocked and Random phases (Tables 3 & 4).

**Fig. 8.**
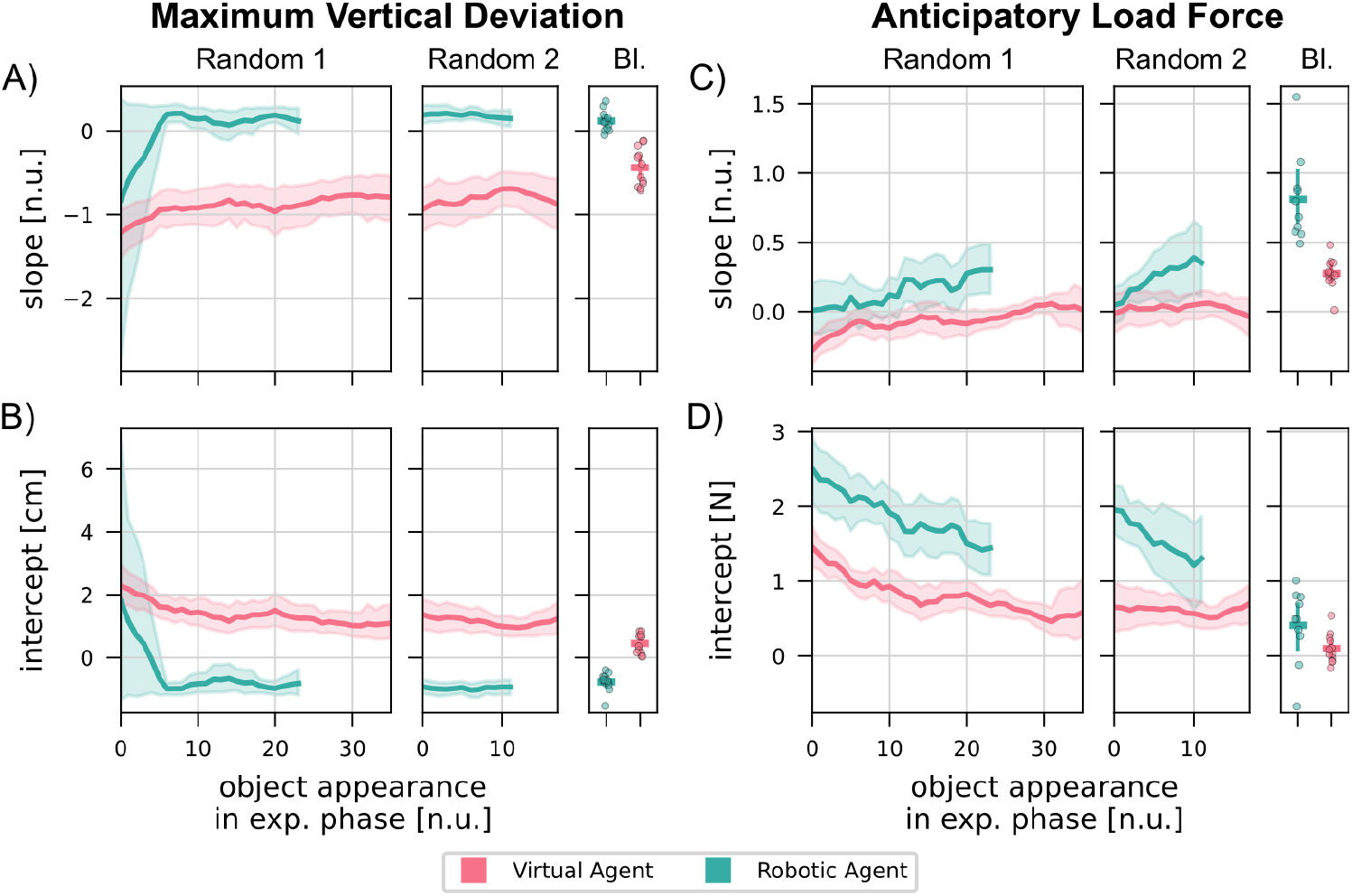
Smoothed trial-by-trial slope and intercept values for all experimental phases in Experiments 1 and 2. A & B) show slope and intercept values for maximum vertical deviation in Virtual and Robotic Agent Experiment, respectively. C & D) show anticipatory load force values in Virtual and Robotic Agent Experiment, respectively. The lines show the mean across participants and shaded area the 95% confidence interval. In the blocked phase of the experiment, the line marker and error bars correspond to the mean and 95% confidence interval across participants, and circular markers correspond to individual participant data.

#### 5.4.2 Anticipatory Load Force

In contrast to all other behavioural metrics, we observe a clear difference between the Blocked and random experimental phases. Within the Blocked phase we observe clear scaling to object mass (Fig.8C; Table3, Table4 Blocked) and a reduced intercept (Fig.8D), this trend does not hold for the random phases of the experiment (Fig.8C; Table3, Table4 Random 1). This pattern is consistent across both experiments. In both experiments, the anticipatory load force slope revealed little scaling to object mass in the random phases of the experiment (see Fig.8C). While the intercept values show a slight decrease over trials, there was a visible offset between the two experiments, with higher intercept values in the robotic agent experiment, indicating an overall higher anticipatory load force (Fig.8D). While steady-state was detected at trial 6 for Virtual Agent Experiment, there was no convergence of the method for Experiment 2.

## 6 Discussion

We examined the effect of different trajectories of the passer in a simulated and physical robot-human object handover on the prediction of object mass of the human receiver. We first demonstrated that our task is valid to investigate prediction of object mass in a handover task. Secondly, we showed that, when objects are presented in a random order, participants were able to understand the kinematic cue and gradually learn to scale their grip forces to object mass in a predictive manner. Finally, we showed that this concept translates from a virtual reality setting to interaction with a physical robot.

### 6.1 Isolating Human Motor Prediction in an Object Handover

We carefully designed two experiments that isolate the participant’s prediction of object mass in a agent-human and robot-human handover. Our handover task is designed such that participants are required to apply a grip force to the object while – held by the passer – it is still locked in space, enabling us to measure purely predictive forces. This is because, prior to the passer releasing the object, no haptic feedback about the object’s mass is available. Once the passer has released the object to the receiver, we can measure the grip forces that these participants would produce given the actual weight, which allows us to compare the predictive grip forces for an object with the corrected grip forces based on full feedback. After a short task familiarization phase, we presented participants with novel object masses in a random order. In this phase we could examine the adaptation of the predictive grip forces purely based on the passer’s kinematic cues. Finally, participants were exposed to the same object masses, but in a blocked fashion. That is, the same mass was demonstrated repetitively, allowing participants to utilize different adaptation mechanisms. The forces observed in this experimental phase can be regarded as the best possible adaptation to a respective object mass.

Using our approach, we demonstrated that when handed an object, participants adapt interaction forces to the object mass on a trial-by-trial basis. While this is expected when repetitively interacting with the same object mass based on error-based adaptation mechanisms [22, 39, 43], we show that even when presented with different object masses in a random order, participants understood the agent’s/robot’s kinematic cue and adapted their grip forces accordingly. Consequently, both the task and experimental design may be used to evaluate legibility and interpretability of robotic cues in future work.

### 6.2 Kinematic Cues to Predict Object Mass

The anticipatory grip force, that is, the grip force applied by the participants before the object was released, provides us with a clear indication of the expected object mass. Therefore, the more closely the anticipatory grip force scales to object mass, the better the prediction accuracy. As we isolated the passing agent’s movement as the only cue in random phases of the experiment, we clearly demonstrate that humans are able to understand this cue and scale their grip forces accordingly. This clearly indicates the formation of a motor memory associating object properties to a kinematic cue. Previous work has shown that visual [7, 29, 46] and even haptic [5, 28] object properties can help us to form such memories. However, we are the first to isolate kinematic cues on the object level and show that interaction dynamics of an agent with the object can trigger similar mechanisms.

Comparing our results to previous work in human-human object handovers, we confirm that a passer kinematic cue at the object level allows the receiver to scale their grip forces in a predictive manner. While previous work hypothesized that observing the passer’s movement is used to predict object mass in a handover [42], we isolate the cue and were able to show that it transfers to non-human passers. In contrast to Kopnarski et al.’s study, we do not observe immediate scaling, but iterative trial-by-trial adaptation evident in iterative increase in slope and decrease in intercept values. As it has been shown, that humans may behave differently when collaborating with robots [8, 40, 53] compared to humans, this is not surprising. While we acknowledge the possibility that robotic movement is treated as a completely arbitrary cue in our experiments, we mainly attribute the increased adaptation to a general lack of familiarity in physical robot-human interaction. Further studies, comparing kinematic to other cues, such as object color or size, are required to fully separate between adaptation to the cue and the task.

To evaluate the strength of the our kinematic contextual cue, we compared the slope and intercept values calculated across all presented object masses for Random and Blocked experimental phases. In the Blocked phase of the experiment, participants were able to use error-driven adaptation mechanisms [22, 39, 43]. That is, they could adapt motor commands based on the errors observed from the previous trial. In contrast, the Random phase of the experiment required context-driven adaptation mechanisms [34, 36].That is, participants had to use the contextual cue, given by the passers movement. In both experiments, we observe a slightly higher slope and lower intercept value in the Blocked phases. This difference between phases is to be expected, due to a large memory effect of the preceding trial on predictive force scaling, as shown in other studies [50].

While a majority of between mass comparisons showed significance of grip force scaling, in the random presentation phase we did not find a significant difference between the 200g and 300g objects in the virtual agent experiment and between the 155g and 255g objects in the robotic agent experiment. Comparable studies investigating human-human handovers in the past commonly used much larger differences in object mass, e.g. [42] with masses ranging from 400g to 1000g. In contrast to these larger differences, the cost of upscaling grip forces by 1N is much lower and, therefore, less likely. Another possibility is, that the scaled component (here: slope) of grip force in our case is “masked” by an increased safety margin related to interacting with another agent. Again, this effect would be expected to be amplified for lower object masses. Finally, especially for the robotic agent experiment, where the handover task took place in the real world, we cannot exclude that the high approach velocity of the robot affected the perceived safety, as observed e.g. in [49], which may have led to an increased baseline grip force, particularly in the low range.

In addition to the anticipatory grip force scaling, we examined the adjustment of grip force after the object was released. The difference between anticipatory and static grip force – the grip force correction – indicates how well the predicted and true object properties matched. This corrective grip force was below 1N in the virtual agent experiment and below 0.5N in the robotic agent experiment, indicating a minimal difference between the initial prediction and the adjusted grip forces after experiencing the object mass. Interestingly, while the adjustments in the real-world experiment were scaled around zero, with a small negative force correction for the light object and a small positive force correction for the heavy object, participants in the virtual reality always showed a positive corrective grip force. That is, for each object mass the static grip force showed an increase compared to the anticipatory grip force. In previous work [32], we observed an increased static grip force intercept when manipulating objects in our VR in contrast to real world experiments with identical mass [59]. We assume that similar mechanisms may have led to the offset of about 0.5N between the two experiments.

In addition to grip force measurements, we measured the applied load force just before the physical handover occurred and the maximum vertical deviation of the object after the release of the object by the passer. Generally, the grip and load force control in object manipulation are tightly coupled [19]. We found that while the anticipatory grip force clearly scaled with the object mass after an initial adaptation period, this was not the case for the anticipatory load force when objects were presented in a random order. Previous studies have reported this phenomenon and attributed this mismatch to be caused by different controllers for the two forces and different task demands [9, 12, 31]. Interestingly, it is not reflected in the recorded vertical object deviation which is reduced from initial to later trials in the Random 1 phases despite the lack of scaling in load forces. The minimal deviation may be the result scaling up load forces in synchrony with the passer’s release. Alternatively, co-contraction of muscles in the shoulder and upper arm and resulting increase in joint stiffness could be a potential strategy to keep the object stationary after the release [23].

Overall, we clearly demonstrate that our participants predictively scale their grip forces to the object mass as cued by the agent/robotic movement. This scaling is learned over multiple trials and occurs despite the visually identical object.

### 6.3 Transfer from Virtual Reality to Real World Human-Robot Interaction

Previous work in robot-human handovers has investigated the effect of biomimicry in both the robotic approach to the handover and the release strategy. Controzzi and colleagues [11] found that the passer release strategy in human-human handover depends on the reaching movement of the receiver and tested this biomimetic strategy with a stationary robotic hand releasing an object to a human participant. In their follow-up work [54], this was expanded to a robot-human handover that included robot reaching movement and a modulation of release strategy according to the human receiver’s reaching movement. However, all experiments were completed with the same object, therefore, after initial familiarization, humans likely formed an accurate prediction about the object’s properties. While we currently focus enabling prediction of the object’s properties, a combination of the two approaches could lead to a more fluent interaction by facilitating prediction of the object’s properties and the partner’s actions. Previous studies have shown that the shape of robot movement trajectory may affect legibility in a handover task [37, 56], with minimum jerk trajectories mimicking human movement emerging as the most predictable for human partners. While we do not investigate different trajectory shapes, we apply this knowledge by using a biomimetic trajectory. To the best of our knowledge, we are the first to show that a manipulation of the trajectory peak velocity can serve as a interpretable contextual cue about object mass.

We first tested our experimental design and kinematic cue using a custom-built virtual reality task. When comparing between the two experiments, we found that while the convergence value of anticipatory grip force slopes was similar, both the intercept convergence values and the overall adaptation was faster in the real-world compared to the virtual agent experiment. Both of these observations are in line with a previous validation study with the same virtual reality setup [32]. Consequently, we attribute the differences between experiments to the uncertainty associated with the overall virtual environment. However, based on our current experiments, we cannot exclude an effect of the difference between a cursor-represented agent and a physical robot. In general, our results validate our virtual reality setup as a testbed for future work related to robot-human object handovers.

## 7 Conclusion and Future Work

Our work demonstrates the possibility to use robot kinematic cues to enable human predictive capabilities in an object handover. We address an important gap in the field of physical human-robot interaction and provide the basis for future work that will allow a seamless and efficient physical collaboration. While this work evaluates important conditions for human predictions solely based on robot movement, we see the potential for future studies evaluating human cognitive load and stress in the field of ergonomics, the strength of a kinematic contextual cue in comparison with other cues and the mechanisms of separate grip and load force control in the field of neuroscience, and the efficiency of physical handover actions using biomimetic release patterns (e.g. [11, 54]) in the field of robotics.

Overall, we validate a task and experiment protocol to measure human prediction in a robot-human handover, and, using this design, demonstrate that robot kinematic cues can be interpreted by human collaborators to predictively scale grasp forces to an object’s mass. By testing our approach both in virtual reality and the real-world, we show generalization across domains.

## Acknowledgments

This work is supported by the Lighthouse Initiatives Geriatronics (Project X, grant no. IUK-1807-0007//IUK582/001) and KI.FABRIK, (Phase 1: Infrastructure as well as the research and development program under grant no. DIK0249) by the Bavarian State Ministry for Economic Affairs, Regional Development and Energy (StMWi), the Deutsche Forschungsgemeinschaft under grant no. 467042759, as well as the TUM Integrative Research Fund, provided by the seed funding initiative of the Munich Institute of Robotics and Machine Intelligence (MIRMI).

## Notes

### Competing Interest Statement

The authors have declared no competing interest.

